# PAN: Personalized Annotation-based Networks for the Prediction of Breast Cancer Relapse

**DOI:** 10.1101/534628

**Authors:** Thin Nguyen, Samuel C. Lee, Thomas P. Quinn, Buu Truong, Xiaomei Li, Truyen Tran, Svetha Venkatesh, Thuc Duy Le

**Affiliations:** Applied Artificial Intelligence Institute, Deakin University, Australia; School of Information Technology and Mathematical Sciences, University of South Australia, Australia

**Keywords:** Personalized medicine, Annotation-based networks, Gene expression, Breast cancer

## Abstract

The classification of clinical samples based on gene expression data is an important part of precision medicine. However, it has proved difficult to accurately predict survival outcomes and treatment responses for cancer patients. In this manuscript, we show how transforming gene expression data into a set of personalized (sample-specific) networks can allow us to harness existing graph-based methods to improve classifier performance. Existing approaches to personalized gene networks all have the limitation that they depend on other samples in the data and must get re-computed whenever a new sample is introduced. Here, we propose a novel method, called Personalized Annotation-based Networks (PAN), that avoids this limitation by using curated annotation databases to transform gene expression data into a graph. These databases organize genes into overlapping gene sets, called annotations, that we use to build a network where nodes represent functional terms and edges represent the similarity between them. Unlike competing methods, PANs are calculated for each sample independent of the population, making it a more efficient way to obtain single-sample networks. Using three breast cancer datasets as a case study (METABRIC and a super-set of GEO studies), we show that PAN classifiers not only predict cancer relapse better than gene features alone, but also outperform PPI and population-level graph-based classifiers. This work demonstrates the practical advantages of graph-based classification for high-dimensional genomic data, while offering a new approach to making sample-specific networks.

**Supplementary information:** The codes and data are available at https://github.com/thinng/PAN.

## 1 Introduction

Breast cancer is one of the leading causes of death for women worldwide, with the incidence and mortality increasing globally [1]. Yet, it is not a single disease, but rather a collection of multiple biological entities, each with their own molecular signature and clinical implications [2]. Since prognosis and treatment response differ between and within cancer sub-types [3], there exists a strong motivation to develop methods that can accurately predict key outcomes for an individual patient, such as relapse after treatment [4]. However, despite advancements in the field, t here r emains much unexplained inter-tumour heterogeneity that drives substantial differences in survival outcomes and treatment responses [5]. Gene expression signatures, easily measured from a tissue sample using high-throughput assays, have been used as a means of stratifying breast cancer samples. This has resulted in computational methods that identify personalized “driver mutation” genes [6], differentially expressed genes and pathways [7], and individualized gene networks [8]. Although genes have been used successfully as biomarkers for cancer prediction tasks [9], it is not clear that gene biomarkers are the most appropriate substrate for classification. Rather, it may be more meaningful to describe diseases in terms of the dysfunction of specific systems, rather than the dysfunction of individual molecules [10]. This perspective is reflected in the use of gene regulatory networks.

Gene regulatory networks represent genes as nodes and the interactions between them as edges. The interactions between genes can be inferred in three ways: from knowledge-driven methods, data-driven methods, or hybrid methods. Knowledge-driven methods use public databases which catalog experimentally confirmed (or predicted) information relating protein-protein interactions (**PPI**) or functionally associated gene sets (called *annotations*). Examples of these databases include the Kyoto Encyclopedia of Genes and Genomes (**KEGG**) [11], Human Phenotype Ontology (**HPO**) [12], Disease Ontology (**DO**) [13], HIPPIE v2.0 [14], among others [15, 16]. Data-driven methods are usually based on gene-gene correlation coefficients [17] or causal relationships [18], as inferred directly from gene expression data. Hybrid methods construct gene regulatory networks by combining gene expression data with the prior knowledge found in annotation databases [19]. Regardless of the method used, most studies compute gene networks at the population- or group-level instead of building personalized (sample-specific) gene regulatory networks. In principle, this results in a single model for all samples in a population, taking a “one-size-fits-all” approach that ignores the inter-tumour heterogeneity of breast cancer. Although population-level networks can help researchers understand a disease in the general sense, personalized gene regulatory networks could pave the way toward accurate and individualized disease prediction.

It is challenging to create sample-specific gene regulatory networks because individuals rarely have the multiple gene expression profiles necessary to compute intra-sample correlations (although single-cell sequencing may offer one way forward [20]). Instead, investigators use custom algorithms to create a sample-specific network. Borgwardt et al. [21] proposed an approach that uses a common PPI reference network to serve as a template from which edges are trimmed based on the gene co-expression status for *that individual* (relative to the population as a whole). Since this method trims edges by comparing a single sample with the population distribution, these PPI-based networks must be re-computed whenever a new sample is introduced. More recently, Kuijjer et al. [22] proposed a more formal method called LIONESS that builds a sample-specific network by estimating the contribution that each sample makes toward the population-level gene regulatory network. As such, LIONESS produces a unique graph for each sample without having to integrate external information. However, LIONESS has the same limitation: whenever a new sample is introduced into the dataset, all sample-specific networks must get r e-computed. Liu et a l. [10] have proposed another method, similar to LIONESS, which also has the same re-computation issue.

In this manuscript, we propose a novel method for constructing sample-specific networks, called Personalized Annotation-based Networks (**PAN**), that uses curated annotation databases to transform gene expression data into a graph. Using the gene set annotations from these databases, we build sample-specific networks where nodes represent functional terms and edges represent the similarity between them. Unlike the PPI-based and LIONESS sample-specific graphs, PAN is calculated for each sample agnostic to the population, removing the need of re-computation, making it a more efficient way to obtain single-sample networks. Using three large breast cancer datasets, we show that the graph properties of PAN not only predict breast cancer relapse data better than gene features alone, but also outperform the PPI-based and LIONESS graphs. To evaluate our method, we test PAN with three separate annotation databases, and find that Disease Ontology (DO) and Human Phenotype Ontology (HPO) databases yield the best performance. This work demonstrates the practical advantages of graph-based classification f or h igh-dimensional g enomic d ata, w hile o ffering a n ew a pproach to making sample-specific networks.

## 2 Methods

### 2.1 Overall objective

The ultimate goal of this work is to predict breast cancer relapse by using personalized (sample-specific) gene regulatory networks for c lassification. For th is, we propose a new method for building sample-specific n etworks, c alled P ersonalized Annotation-based Networks (**PAN**), that we benchmark against established methods. Since PAN depends on the annotation database chosen, we test three databases: Kyoto Encyclopedia of Genes and Genomes (**KEGG**) [11], Human Phenotype Ontology (**HPO**) [12], and Disease Ontology (**DO**). To evaluate the performance of PAN, we use publicly available data from the Gene Expression Omnibus (**GEO**) and METABRIC. Our proposed PAN method is summarized in Figure 1.

**Figure 1:**
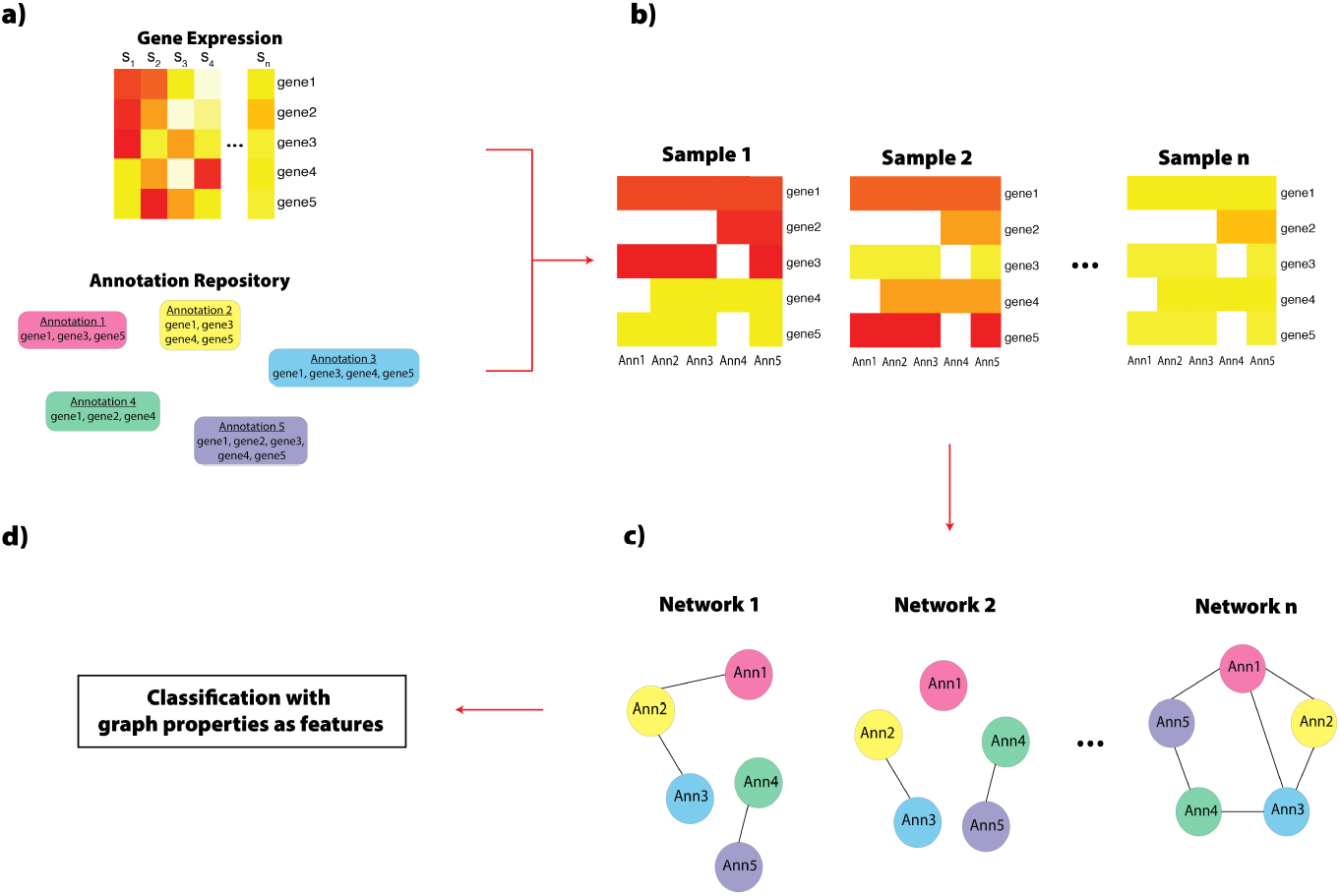
This figure illustrates how we create a Personalized Annotation-based Network (**PAN**) graph. **Panel A:** We obtain publicly available Gene Expression datasets (GEO-5 or METABRIC) and Gene Annotations (KEGG, HPO, or DO). **Panel B:** For each sample, we build an intermediate matrix by matching the Gene Expression data to the Gene Annotations: if there is an annotation for that gene, the value of the intermediate matrix becomes the expression of that gene. Otherwise, the value becomes zero. **Panel C:** From the intermediate matrix, we build a network where the nodes are the annotations and the edges are the Euclidean distance between them. To turn the annotation-annotation association weights into a discretized network, we create an adjacency matrix by selecting the top 10% of edges with the smallest Euclidean distance. **Panel D:** After building the network, we use its properties (Betweenness Centrality, Closeness Centrality, or PageRank) as feature input for a classification algorithm to predict breast cancer relapse. Acronyms: S (Sample); Ann (Annotation).

### 2.2 Data acquisition

The data used for the prediction of events were obtained from two main sources, the Gene Expression Omnibus (**GEO**) and METABRIC. The first dataset, named **GEO-5**, is a super-set of five GEO collections (including GSE12276 [23], GSE20711 [24], GSE19615 [25], GSE21653 [26], and GSE9195 [27]). These datasets were selected because they all use the same micro-array platform (Affymetrix Human Genome U133 Plus 2.0 Array) and include clinical information about relapse. The GEO-5 dataset contains 736 samples (349 relapse; 387 no relapse).

The second dataset was retrieved from METABRIC, a popular breast cancer dataset from the European Genome-Phenome Archive [28]. We obtained the gene expression data already pre-processed by the limma [29] and fRMA [30] libraries. Next, we adjusted the data for batch effects using the ComBat algorithm from the sva library [31, 32] (with default parameters and covariates as tumor versus normal). The repository contains 2000 breast tumor samples. To be included in our study, we required that the sample had clinical information about relapse. This reduced the dataset to 1283 samples (422 relapse; 861 no relapse).

The third dataset, named **UK207**, was compiled from the GSE22216 and GSE22220 collections, both of which used the Illumina humanRef-8 v1.0 beadchip expression array [33]. We retrieved the data already log2-normalized for 210 early primary breast cancer samples taken during a 10-year follow-up. These patients underwent various treatments and were evaluated for relapse. After removing samples with missing information in the survival meta-data, 207 samples remained (77 relapse; 130 no relapse).

### 2.3 Defining sample-specific networks

This section describes three sample-specific networks experimented in this work: our proposed Personalized Annotation-based Networks and two baselines: PPI-based and LIONESS.

#### 2.3.1 PPI-based networks

In this method, a protein-protein interaction (**PPI**) network [21] serves as reference network that is subsequently pruned to establish sample-specific networks. For each sample, we keep any PPI edge where its constituent genes both have high (or low) relative expression. Specifically, we include all edges where both genes are in either the top (or bottom) quantiles as compared with the other samples. For this procedure, we use the HIPPIE database of experimentally observed PPIs [14].

#### 2.3.2 LIONESS networks

We applied LIONESS to the gene expression data using PyPanda [34], run with default parameters.

#### 2.3.3 Personalized Annotation-based Networks (PAN)

In this work, we propose a novel method, called Personalized Annotation-based Networks (**PAN**), to build personalized networks for individual samples based on gene expression data. We begin by using an annotation database (KEGG, HPO, or DO) to convert gene expression data into an annotation-based network. This is done by calculating the intra-sample distance between pairs of annotations, the magnitude of which becomes the edge weight. Formally, each sample *j* is represented by the matrix 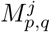, where *p* is a gene and *q* is an annotation (which may or may not be associated with gene *p*). The value for each element 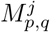 equals the expression of gene *p* if it is associated with the annotation *q*. Otherwise, it equals zero.

Then, M^*j*^ is turned into a symmetric association matrix, A^*j*^, where the value for each element 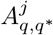 describes the Euclidean distance between any two annotations *q* and *q*^*^. For PAN, the nodes are annotations and the edges are the Euclidean distance between them. Since the distance between two annotations will equal 0 when both sets have the same genes, A^*j*^ effectively measures the total abundance of all genes present in only one of the two sets. To turn the annotation-annotation association weights into a discretized network, we create an adjacency matrix from the top 10% of edges as ranked by association strength. Since a network is created for each sample independently, PAN does not have the limitation that the PPI-based and LIONESS approaches have, and can thus work efficiently for streaming data. Our proposed PAN method is summarized in Figure 1.

### 2.4 Graph-based Representation

Once the discretized networks are constructed for each sample, their graph properties can be extracted and used as features for classification. Given a sample *S*, a graph *G^S^ V^S^, E^S^* is constructed from its gene expression using the PPI-based method [21], the LIONESS method [22], or the proposed PAN method. The graph-based representation of *S* is denoted as **h**^*S*^ and defined based on *G^S^*. In the following sub-sections, we present different ways to define **h**^*S*^. By using graph properties, **h**^*S*^ is represented as a vector of V^*S*^ dimensions, i.e., 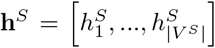 where 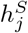 is computed from properties of the vertex 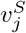.

#### 2.4.1 Closeness Centrality

The Closeness Centrality of a node 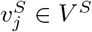 is defined as the reciprocal of the sum of the shortest path distances from 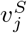 to all other nodes [35],

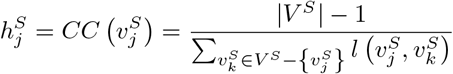

where 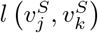 is the length of the shortest-path from 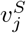 to 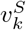.

#### 2.4.2 Betweenness Centrality

The Betweenness Centrality of a node 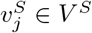 is defined as the sum of the fraction of all-pairs shortest paths that pass through 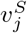 [36],

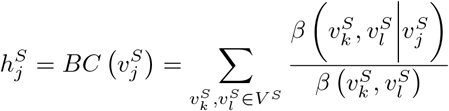

where 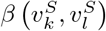 is the number of shortest 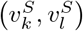-paths and 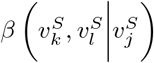 is the number of those paths passing through 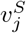 other than 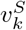 and 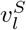. Note if 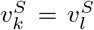, then 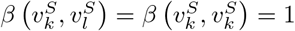, and if 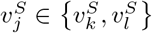, then 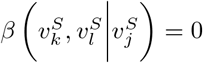.

**Table 1:** This table shows the maximum average cross-validation AUC observed for each classification method and dataset studied, regardless of the number of genes or classifier model (LR vs. SVM) used

| Dataset | LIONESS | non-graph | PAN_DO | PAN_HPO | PAN_KEGG | PPI-based |
| --- | --- | --- | --- | --- | --- | --- |
| GEO-5 | 0.5671 | 0.5293 | **0.6473** | 0.6418 | 0.5804 | 0.6443 |
| METABRIC | 0.6004 | 0.6176 | **0.6262** | 0.6225 | 0.6259 | 0.5596 |
| UK207 | 0.6504 | 0.6488 | 0.6800 | **0.6832** | 0.7214 | 0.6145 |

#### 2.4.3 PageRank

PageRank [37] was developed for measuring the importance of websites on the Internet. This method is based on an underlying assumption that the most important websites are more likely to receive links from other sites. In our case, we define the PageRank property of a node 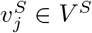 as:

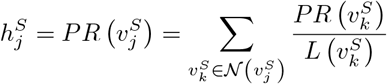

where 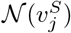 is the set of all nodes linking to node 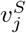 and 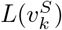 is the number of links from 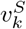.

### 2.5 Classifier choice and performance

We trained logistic regression (**LR**) and linear kernel support vector machine (**SVM**) [38] classifiers using (a) the gene expression data (**non-graph**) and (b) the properties of the PPI-based, LIONESS, and PAN graphs. Since PAN depends on the database used, we trialled three databases, resulting in three separate PAN models: **PAN_KEGG**, **PAN_HPO**, and **PAN_DO**. For each classifier, we report all performance measures as averaged across 10 test sets (i.e., using bootstrapped cross-validation, also known as Monte Carlo cross-validation).

## 3 Results and Discussion

### 3.1 Sample-specific networks predict breast cancer relapse

Once a single-sample network is created, its graph properties can used as feature input for the prediction of breast cancer relapse. Figure 2 shows the average cross-validation AUC for graph and non-graph classifiers. This figure compares PAN classifier performance with the PPI, LIONESS, and non-graph classifiers. For the GEO-5 dataset, we see that all graph classifiers perform better than the commonly used non-graph (i.e., gene expression) classifier. Considering that the GEO-5 dataset is aggregated from multiple sources, this might suggest that single-sample networks capture a signal that is more robust to inter-batch differences. For the other datasets, the PPI-based and LIONESS graphs do not perform as well, although the PAN graphs still have among the best performance. Table 2.5 summarizes the figure by showing the single best average cross-validation AUC regardless of the number of genes or classifier model used.

**Table 2:**
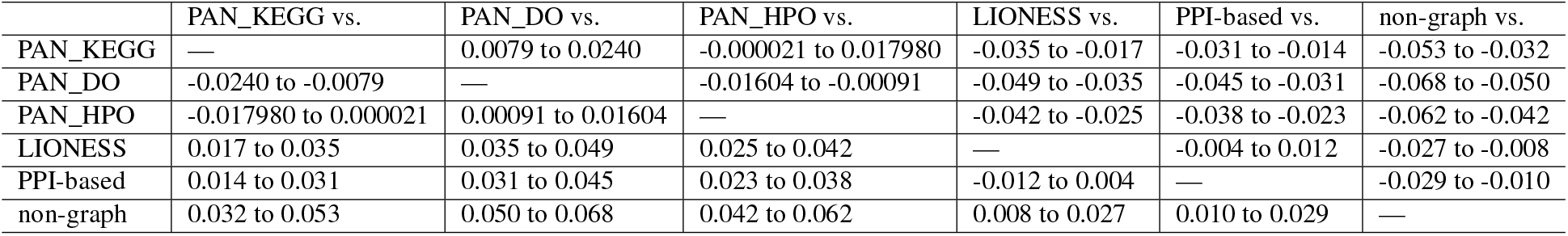
This table shows the 95% confidence interval for the median of the differences between logistic regression (LR) classifiers, computed using pair-wise Wilcoxon Rank-Sum tests for all classifier sizes, datasets, and cross-validation folds. Here, we see that the PAN_DO method outperforms all other methods, having at least 3% and 5% better AUC than the competing graph and non-graph methods, respectively.

**Table 3:**
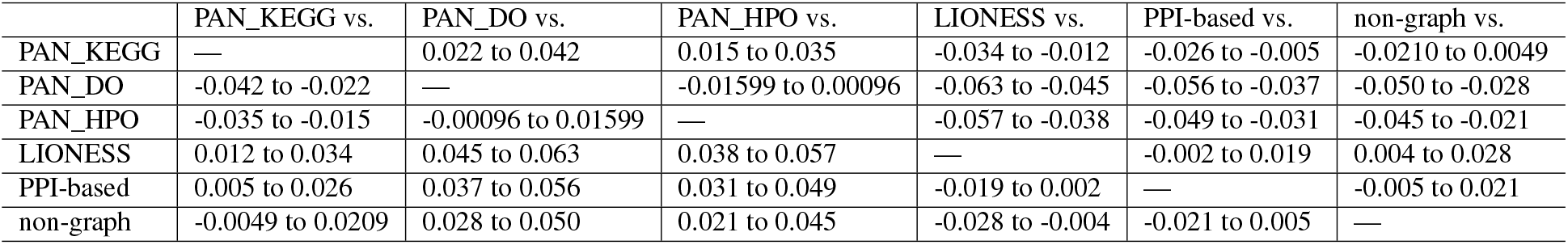
This table shows the 95% confidence interval for the median of the differences between linear kernel support vector machine (SVM) classifiers, computed using pair-wise Wilcoxon Rank-Sum tests for all classifier sizes, datasets, and cross-validation folds. Here, we see that the PAN_DO and PAN_HPO methods perform best, having at least 3% and 2% better AUC than the competing graph and non-graph methods, respectively.

|  | PAN_KEGG vs. | PAN_DO vs. | PAN_HPO vs. | LIONESS vs. | PPI-based vs. | non-graph vs. |
| --- | --- | --- | --- | --- | --- | --- |
| PAN_KEGG | — | 0.022 to 0.042 | 0.015 to 0.035 | -0.034 to -0.012 | -0.026 to -0.005 | -0.0210 to 0.0049 |
| PAN_DO | -0.042 to -0.022 | — | -0.01599 to 0.00096 | -0.063 to -0.045 | -0.056 to -0.037 | -0.050 to -0.028 |
| PAN_HPO | -0.035 to -0.015 | -0.00096 to 0.01599 | — | -0.057 to -0.038 | -0.049 to -0.031 | -0.045 to -0.021 |
| LIONESS | 0.012 to 0.034 | 0.045 to 0.063 | 0.038 to 0.057 | — | -0.002 to 0.019 | 0.004 to 0.028 |
| PPI-based | 0.005 to 0.026 | 0.037 to 0.056 | 0.031 to 0.049 | -0.019 to 0.002 | — | -0.005 to 0.021 |
| non-graph | -0.0049 to 0.0209 | 0.028 to 0.050 | 0.021 to 0.045 | -0.028 to -0.004 | -0.021 to 0.005 | — |

**Figure 2:**
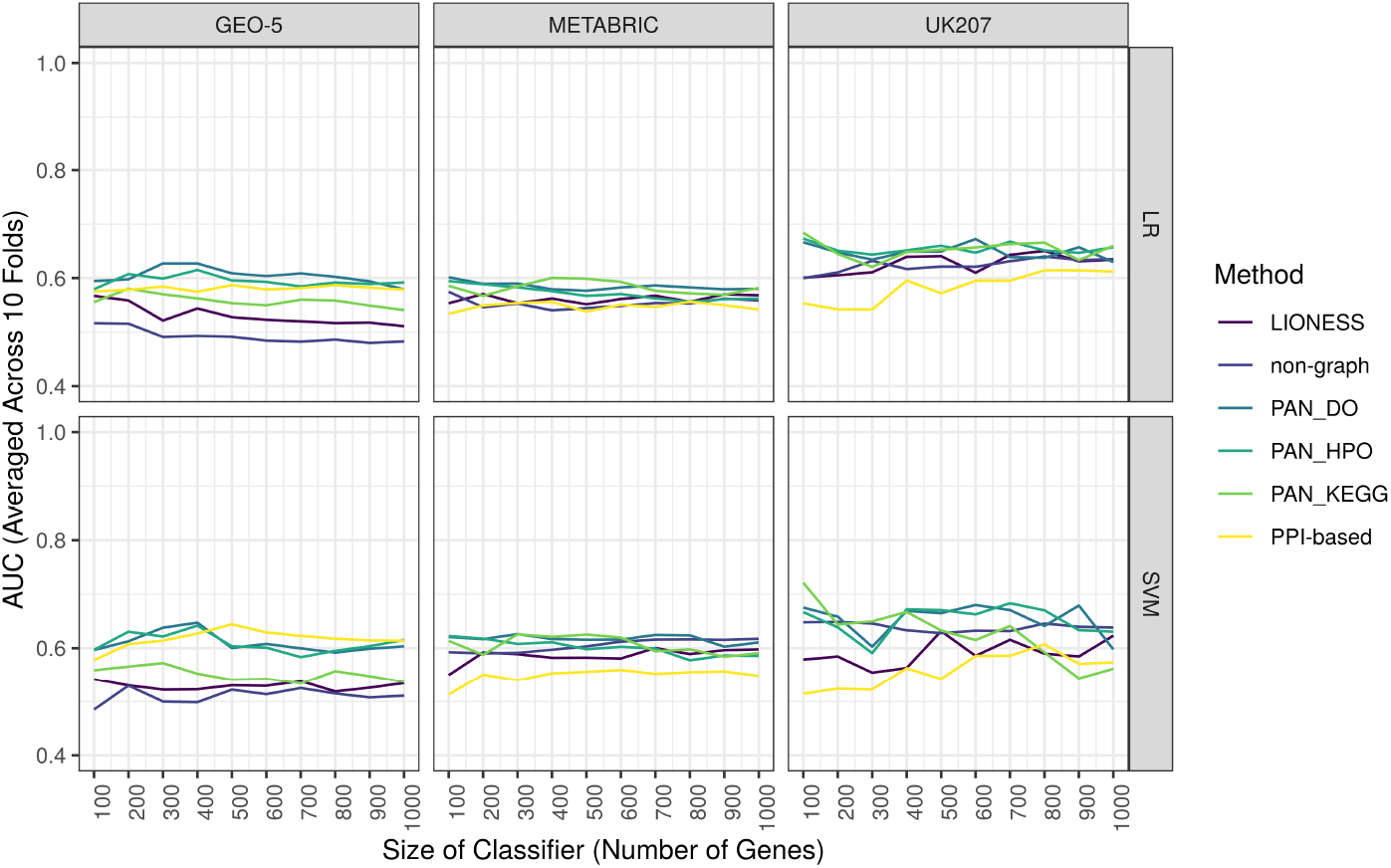
This figure shows the average AUC (y-axis) of graph and non-graph methods (color) trained on the top *G* = [100, …, 1000] most variable genes (x-axis). The x-facet shows the 3 datasets under study, while the y-facet shows the classification algorithm used (LR vs. SVM). PPI, LIONESS, and PAN are graph-based methods, where graph properties are used as feature input. The non-graph method uses gene expression data as feature input. Here, we see how the PAN methods consistently outperform all graph and non-graph methods. LIONESS and non-graph methods tend to perform the worst. All performance metrics averaged across 10 test sets.

Although PAN graphs are sample-specific, we can summarize their architecture across the population in order to visualize their discriminative potential. Figure 3 shows composite graphs for each of the datasets, made using 100 genes and the HPO annotation database. In these figures, an edge indicates that at least 80% of the samples within a group (Relapse or No-Relapse) contain the edge. Here, we can easily see several edges which are present in >80% of Relapse samples but not in No-Relapse samples. The right panel of the figure shows a frequency histogram of the difference in class proportions. Here, we see that some edges are strongly associated with Relapse only (or No-Relapse only), though much intra-class variability exists.

**Figure 3:**
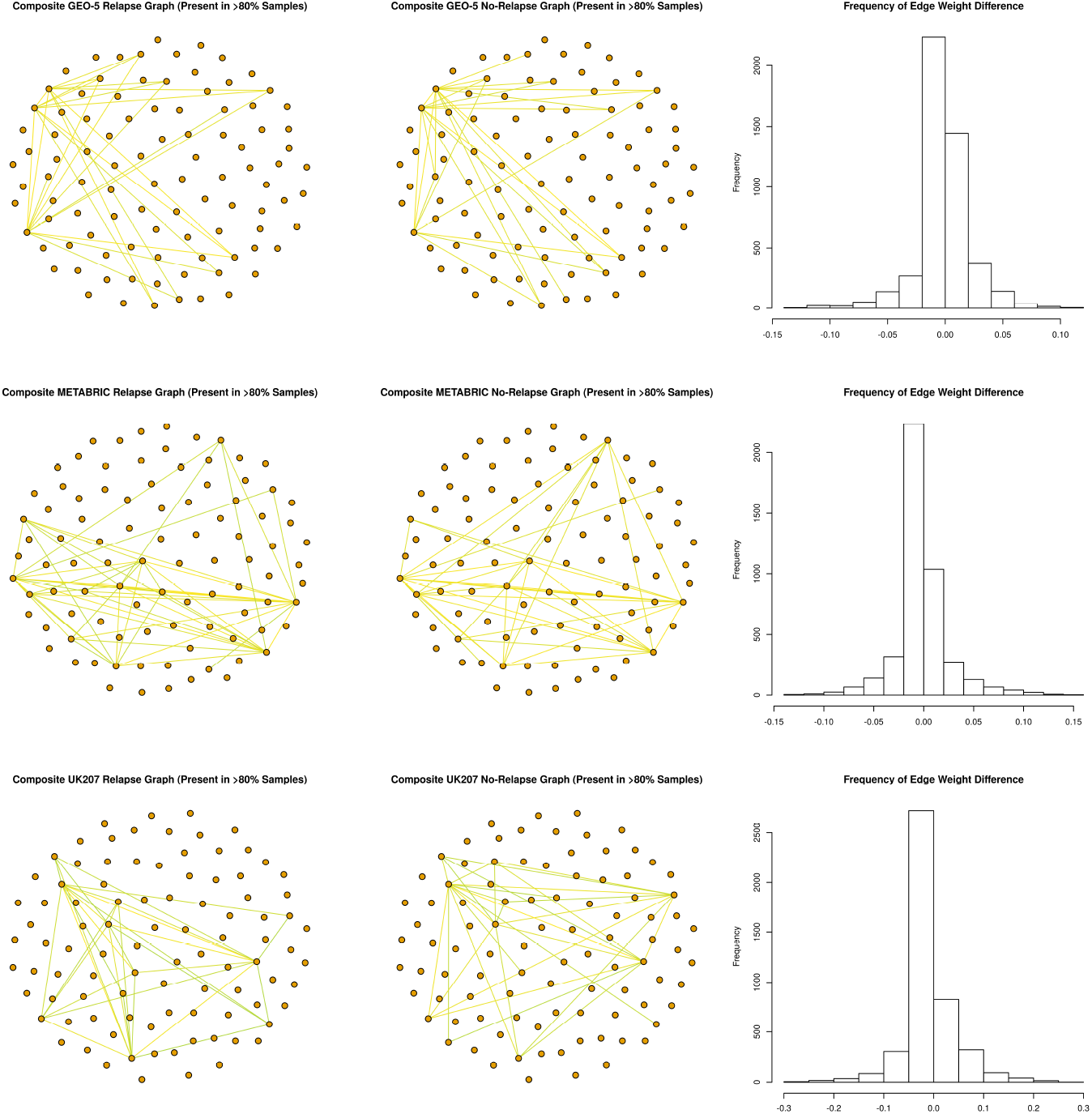
This figure shows composite graphs for the Relapse (left column) and No-Relapse (middle column) samples for each dataset (as row), made using 100 genes and the HPO annotation database. For composite graphs, an edge indicates that at least 80% of the samples contain the edge after filtering. The histogram (right column) shows the frequency of the difference in class proportions, equal to the proportion of the Relapse samples with an edge minus the proportion of No-Relapse samples with the edge. Note that nodes only have a fixed position for a given dataset (as row).

### 3.2 PAN graphs outperform competing methods

The goodness of any classification method might depend on the dataset under study. Therefore, it is important to evaluate a classification method across multiple datasets. When comparing the rank-order of the six methods across 3 datasets, we see that the PPI-based classifier has an inconsistent performance. Although the PPI-based classifier is among the best for GEO-5, it is among the worst for METABRIC (see Figure 2: SVM facet). On the other hand, the LIONESS and non-graph (i.e., gene expression) classifiers consistently have poor or average performance. Meanwhile, the PAN method out-performs all competing methods. Although the margin is small, PAN_DO usually performs better than PAN_HPO, both of which usually perform better than PAN_KEGG. As such, we see a stable rank-order among the PAN methods: PAN_DO > PAN_HPO > PAN_KEGG. Tables 2.5 and 2.5 show the 95% confidence interval for the median of the differences between classification methods, computed across all classifier sizes, datasets, and cross-validation folds. This table shows that PAN_DO can yield a 2-6% better AUC than non-PAN methods. It is interesting to note that the better performing annotation databases have fewer total annotations. This could suggest that PAN’s success may have something to do with its ability to condense the high-dimensional gene expression data into a lower-dimensional space.

## 4 Summary

In this paper, we propose a novel method for constructing sample-specific networks, called Personalized Annotation-based Networks (**PAN**), which use curated annotation databases to transform gene expression data into a graph. Using the properties of these graphs as feature input for classification, we show that PAN not only predicts breast cancer relapse better than non-graph (i.e., gene expression) classifiers, but also outperform competing sample-specific network methods. Although PAN graphs depend on the annotation database used, we show that PAN classifiers perform consistently well for 3 annotation databases (KEGG, HPO, and DO), with PAN_DO having superior performance in most tests. Our results support two principal conclusions. First, they suggest that applying graph-based models to the classification of gene expression data improves performance considerably. Second, they suggest that integrating annotation databases into classification pipelines i s appropriate for clinically relevant classification problems, such as the prediction of breast cancer relapse. Although we showcase the PAN method on gene expression data, our method can be generalized to any classification problem where a relevant annotation database exists.

## Declarations

### Ethics approval and consent to participate

Not applicable.

### Consent to publish

Not applicable.

### Availability of data and material

The PAN model and all baselines are implemented in Python. The code is freely available from https://github.com/thinng/PAN.

## Acknowledgements

Not applicable.

